# beta-blocker reverses inhibition of beta-2 adrenergic receptor resensitization by hypoxia

**DOI:** 10.1101/2020.09.17.301903

**Authors:** Yu Sun, Manveen K. Gupta, Kate Stenson, Maradumane L. Mohan, Nicholas Wanner, Kewal Asosingh, Serpil Erzurum, Sathyamangla V. Naga Prasad

## Abstract

Ischemia/hypoxia is major underlying cause for heart failure and stroke. Although beta-adrenergic receptor (βAR) is phosphorylated in response to hypoxia, less is known about the underlying mechanisms. Hypoxia results in robust GRK2-mediated β2AR phosphorylation but does not cause receptor internalization. However, hypoxia leads to significant endosomal-β2AR phosphorylation accompanied by inhibition of β2AR-associated protein phosphatase 2A (PP2A) activity impairing resensitization. Phosphoinositide 3-kinase γ (PI3Kγ) impedes resensitization by phosphorylating endogenous inhibitor of protein phosphatase 2A, I2PP2A that inhibits PP2A activity. Hypoxia increased PI3Kγ activity leading to significant phosphorylation of I2PP2A resulting in inhibition of PP2A and consequently resensitization. Surprisingly, β-blocker abrogated hypoxia-mediated β2AR phosphorylation instead of phosphorylation in normoxia. Subjecting mice to hypoxia leads to significant cardiac dysfunction and β2AR phosphorylation showing conservation of non-canonical hypoxia-mediated pathway in vivo. These findings provide mechanistic insights on hypoxia-mediated βAR dysfunction which is rescued by β-blocker and will have significant implications in heart failure and stroke.

## Introduction

Oxygen is a key currency driving the sustenance of the cells as it plays a central role in metabolism and respiration [1]. The need of oxygen for such a fundamental and existential physiology has led the eukaryotes to develop exquisite mechanisms to maintain and match the ever changing needs of oxygen by the cells/tissues [1, 2]. The reduction in oxygen supply classically occurs due to increased demand that exceeds the supply either locally or systemically causing hypoxia leading to metabolic crisis with implications in cell survival. In recognition of the critical role oxygen plays in functional homeostasis, eukaryotes have developed an efficient and rapid oxygen sensing system, the hypoxia-inducible factors (HIFs) which are master transcription factors [2–5]. HIF family is represented by members HIF-1, −2 and −3 off which HIF-1α isoform is most well studied. HIF-1α is stabilized in hypoxia and dimerizes with HIF-1β to form a potent transcription factor that drives the hypoxia response [2, 6–8]. However, our previous work has shown that HIF-1α can be stabilized by beta-adrenergic receptor (βAR) activation in normoxia [9]. While, it is also known that β2ARs are regulated by oxygen through hydroxylation that alters β2AR stability and responses [9, 10].

βARs are prototypic G-protein coupled receptors (GPCRs) that play a key role in cardiac function [11] wherein their sustained dysfunction is associated with deleterious cardiac remodeling and heart failure [11, 12]. There are three sub-types of βARs (β1, β2, and β3AR) of which β2AR is ubiquitously expressed while, β1AR is primarily expressed in the heart. Agonist binding to βARs like endogenous ligands epinephrine and norepinephrine leads to G-protein coupling resulting in dissociation of hetero-trimeric G-protein into Gαs and Gβγ subunits, and cAMP generation [11, 13–15]. The dissociated Gβγ subunits recruit GPCR kinase 2 (GRK2) to the receptor leading to βAR phosphorylation and desensitization *ie* inability to couple to G-protein despite agonist [11, 16]. As adaptor scaffolding protein β-arrestin binds to phosphorylated βAR and targets it for endocytosis [17, 18]. The endocytosed β2AR undergoes dephosphorylation in the endosomes before being recycled back to the plasma membrane as naïve receptors [17, 19]. Our previous studies have shown that phosphoinositide 3-kinase γ (PI3Kγ) that is recruited to the receptor complex inhibits protein phosphatase 2A (PP2A) at plasma membrane through phosphorylation of inhibitor of PP2A (I2PP2A) [20]. Thus, agonist activation leads to increased phosphorylation of β2AR due to kinase activity of GRK2 and simultaneous inhibition of PP2A by PI3Kγ. In contrast to this traditional agonist mediated mechanisms, we have shown that hypoxia causes β2AR phosphorylation in the absence of agonist which is associated with HIF-1α accumulation [9]. Furthermore, GRK2 inhibition results in reduction of hypoxia induced-β2AR phosphorylation and HIF1-α accumulation [9]. Interestingly, despite agonist-independent βAR dysfunction, use of β-blocker surprisingly reduces HIF-1α accumulation with hypoxia [9].

Traditionally, β-blockers are antagonists that block the activation of the βARs and as a consequence there is no G-protein coupling and cAMP generation [21]. Accumulating evidence has shown that β-blocker can mediate biased signaling wherein, they block G protein-dependent signaling while simultaneously initiating G protein-independent β-arrestin dependent signaling [22, 23]. Studies have shown that β-blockers mediate downstream G protein-independent signaling through EGFR transactivation [21]. This suggests that β-blockers are able to confer a unique βAR conformation that allows for G protein independent signaling. Consistent with this idea that βARs can attain different conformations that allows receptors to activate unique downstream signal, studies have shown that hypoxia leads to unique phosphorylation bar-code on the receptor that regulates HIF-1α accumulation [9]. Given that the mechanistic underpinnings of this regulation is not well understood, our current studies have focused on identifying the determinants of the unique non-canonical agonist-independent hypoxia mediated regulation of β2AR function. We show in our current study that in addition to selective upregulation of GRK2, there also simultaneous inhibition of PP2A-mediated resensitization accounting for accumulation of phosphorylated β2ARs. Consistent with inhibition of resensitization, there is significant increase in endosomal PI3Kγ activity and concomitant reduction in β2AR-associated phosphatase activity. Moreover, there was marked increase in I2PP2A phosphorylation accounting for the loss in PP2A activity. Correspondingly, subjecting mice to acute hypoxia resulted in deleterious cardiac remodeling associated with significant βAR dysfunction showing conservation of these pathways in vivo. Surprisingly, β-blocker propranolol in hypoxia reversed β2AR phosphorylation in contrast to normoxia wherein, it consistently induced β2AR phosphorylation showing non-canonical regulation of β2ARs by hypoxia.

## Results

### Selective increase in GRK2 mediates hypoxia-indueed β2AR dysfunction

To test whether hypoxia *per se* causes β2AR phosphorylation, HEK 293 cells stably expressing FLAG-β2AR (β2AR-HEK 293 cells) were serum starved and subjected to hypoxia for 0, 3 and 6 hours. Immunoblotting of cell lysates with anti-phospho-β2AR antibody showed significant phosphorylation of β2AR by 6 hours (n=5) [**Fig. 1A**]. The membranes were stripped and reblotted for FLAG as a loading control for the β2AR expression in the cells [**Fig. 1A**]. To further determine whether the cells were subjected to hypoxia stress, the membranes were stripped and re-immunoblotted with anti-HIF-1α antibody. Accumulation of HIF-1α occurred by 6 hours with no appreciable difference at 3 hours of hypoxia treatment [**Fig. 1A**] showing increase in β2AR phosphorylation associated with HIF-1α accumulation. Confocal microscopy showed that in contrast to normoxia, hypoxia resulted in significant accumulation of phosphorylated β2AR as visualized by anti-phospho-β2AR antibody (green) (n=4)[**Fig. 1B (panels 2 & 6) and C**].

**Figure 1.**
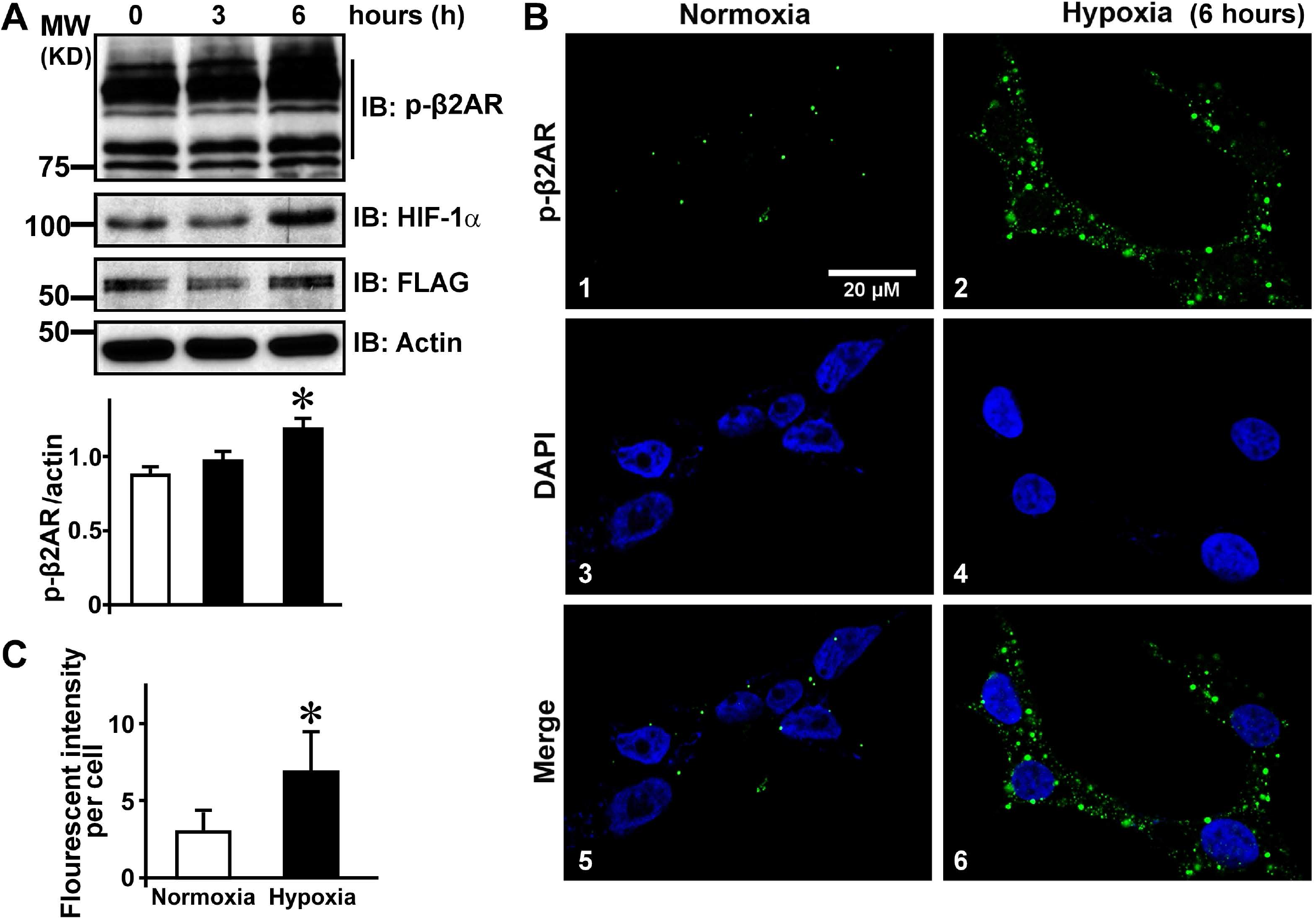
Hypoxia leads to increased phosphorylation of β2AR: **A**, Total cell lysates (80 μg) from serum starved HEK 293 cells stably expressing FLAG-β2AR (β2AR-HEK 293 cells) following 0, 3 and 6 hours (h) of hypoxia (2% oxygen) or normoxia were immunoblotted with anti-phospho-β2AR antibody to assess β2AR phosphorylation (upper panel). The immunoblot was stripped and re-probed with anti-HIF-1α antibody to determine expression of HIF-1α as a molecular surrogate for hypoxia. The immunoblots were probed with anti-FLAG and anti-actin antibody as loading controls. Cumulative densitometric analysis of five independent experiments (n=5), *p< 0.01 vs. 0 and 3 hour time point. **B**, Confocal images of cell stained with anti-phospho-β2AR antibody (green) after 6-hour normoxia or hypoxia treatment. Nucleus was visualized by DAPI (blue) staining. Scale, 20 μm. **C**, Fluorescent intensity of phosphorylated β2ARs/cell. Over 70-100 cells/experiment were used for fluorescent assessment, (n=4), *p< 0.05.

Since GRKs mediate phosphorylation of β2ARs, cell lysates were assessed to determine which GRKs are involved in mediating receptor phosphorylation in response to hypoxia. Comprehensive immunoblotting for ubiquitously expressed GRKs (GRK 2, 3, 5 and 6 [24]) showed selective and significant increase only in GRK2 expression (n=5) [**Fig. 2A & B**] with no appreciable changes in other GRKs [**Fig. 2A**]. Given the consistent correlation between GRK2 upregulation and β2AR phosphorylation at 6 hours, all the studies described from hereon used 6 hours of hypoxia treatment for assessing mechanistic underpinnings of βAR dysfunction. To test whether GRK2 activity is sufficient to mediate β2AR phosphorylation in response to hypoxia, β2AR-HEK 293 cells were pre-treated with GRK2 inhibitor paroxetine. Paroxetine treatment abrogated the hypoxia-mediated phosphorylation of β2ARs (n=4) [**Fig. 2C**] showing the GRK2 is the key kinase that phosphorylates β2ARs following hypoxia. Given that hypoxia causes β2AR phosphorylation indicating receptor dysfunction, immediate downstream signal was assessed by measuring cAMP level in β2AR-HEK 293 cells. Consistent with the loss in β2AR function, there was significant reduction in the amount of cAMP following hypoxia (n=5) [**Fig. 2D**]. To further test whether G protein coupling is altered following hypoxia, plasma membranes and endosomal fractions were isolated from normoxia and hypoxia treated cells. These fractions were subjected to in vitro isoproterenol (ISO) (βAR agonist) stimulation to assess for G protein coupling. Increased adenylyl cyclase activity was observed following ISO stimulation in the plasma membrane (n=5) [**Fig. 2E]** as well as in endosomal fractions (n=5) [**Fig. 2F**] in normoxia treated cells. While significant reduction in adenylyl cyclase activity was observed in hypoxia treated cells showing that hypoxia-induced β2AR phosphorylation impairs receptor activation promoting β2AR dysfunction.

**Figure 2.**
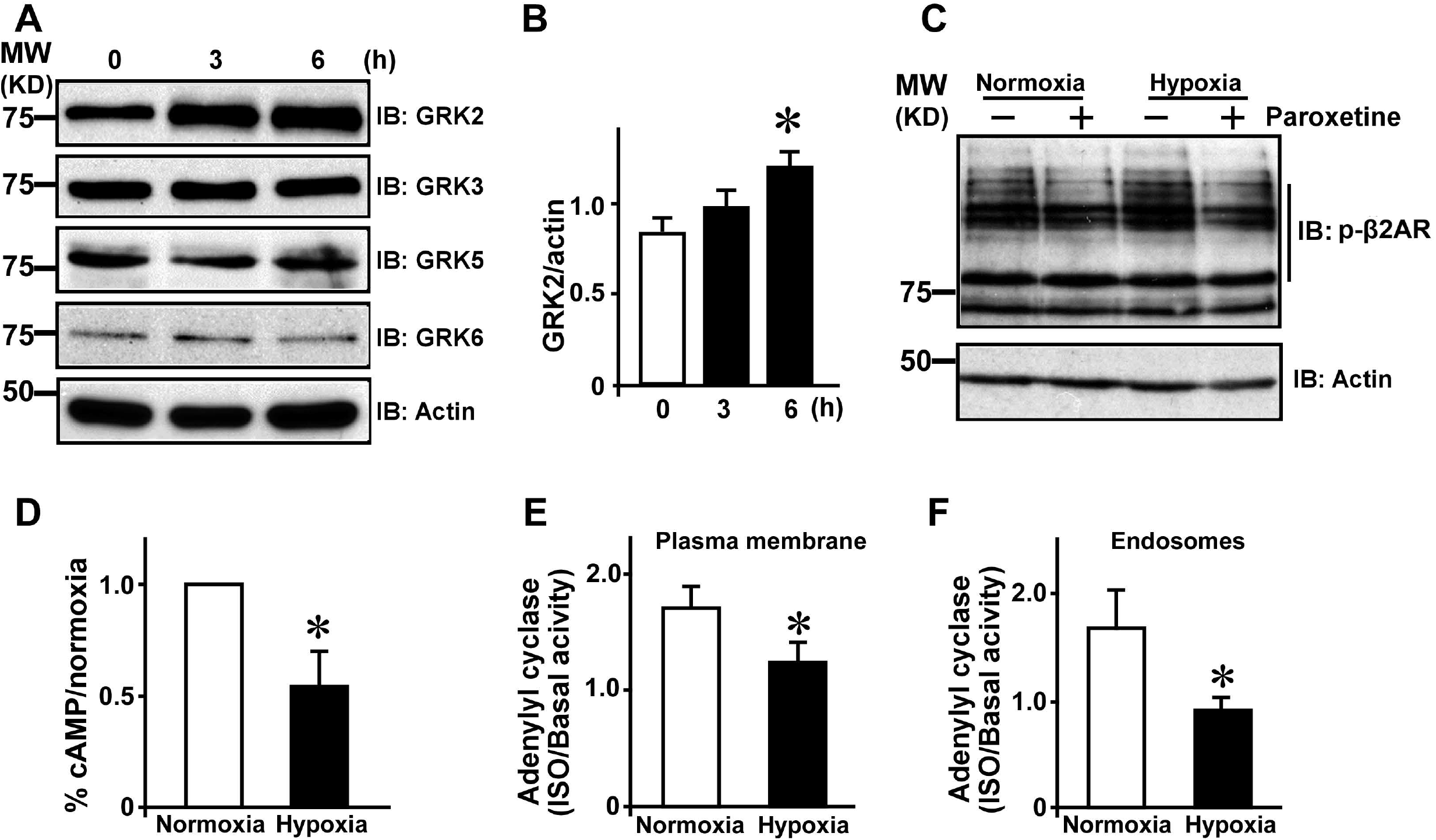
Hypoxia mediates β2AR dysfunction through GRK2: **A**, Total cell lysates (80 μg) were immunoblotted for ubiquitously expressed GRKs, GRK2, 3, 5 or 6 from β2AR-HEK 293 cells following 6 hours of normoxia or hypoxia. **B**, Summary densitometric analysis of GRK2 (n=5), *p<0.05 vs. 0 hour. **C**, β2AR-HEK 293 cells were pre-treated with GRK2 inhibitor (paroxetine, 30μM) 45 minutes prior to 6 hours of hypoxia or normoxia treatment and total cell lysates (80 μg) were immunoblotted with anti-phospho-β2AR antibody. **D**, cAMP levels were measured in β2AR-HEK 293 cells following 6 hours of hypoxia treatment compared to normoxia (n=5). *p<0.001 vs normoxia. **E & F**, In vitro isoproterenol (ISO)-stimulated adenylyl cyclase activity was measured in the plasma membrane (E) and endosomal fractions (F) extracted from β2AR-HEK 293 cells after 6 hours of hypoxia or normoxia. The data is presented as fold change following in vitro ISO-stimulation/baseline (n=5). Plasma membranes (E) *p<0.05 vs normoxia and (F) endosomes, *p<0.01 vs normoxia.

### Hypoxia disengages β2AR phosphorylation-internalization axis

Traditionally, β2AR phosphorylation by GRK2 mediates β-arrestin recruitment leading to desensitization and internalization of the receptors into the endosomes. To test whether β-arrestin plays a role in hypoxia-induced β2AR desensitization and dysfunction, β-arrestin 2 GFP-β2AR double-stable HEK 293 cells underwent normoxia or hypoxia treatment (6 hours) or ISO stimulation for 10 minutes (that was used as a positive control). Consistent with previous studies [25], ISO stimulation resulted in significant recruitment of β-arrestin 2 GFP to the plasma membrane (green) (n=3) [**Fig. 3A, panels 5 & 8**] as assessed by the clearance of the cytosolic β-arrestin 2 GFP. This is associated with a subset of β2ARs that are phosphorylated (red) [**Fig. 3A, panels 6 & 8**] and internalized. While hypoxia results in marked phosphorylation of β2ARs (red) [**Fig. 3A, panels 10 & 12**], no appreciable changes in distribution of β-arrestin 2 GFP was observed [**Fig. 3A, panels 9 & 12**] suggesting non-canonical regulation of β2AR by hypoxia.

**Figure 3.**
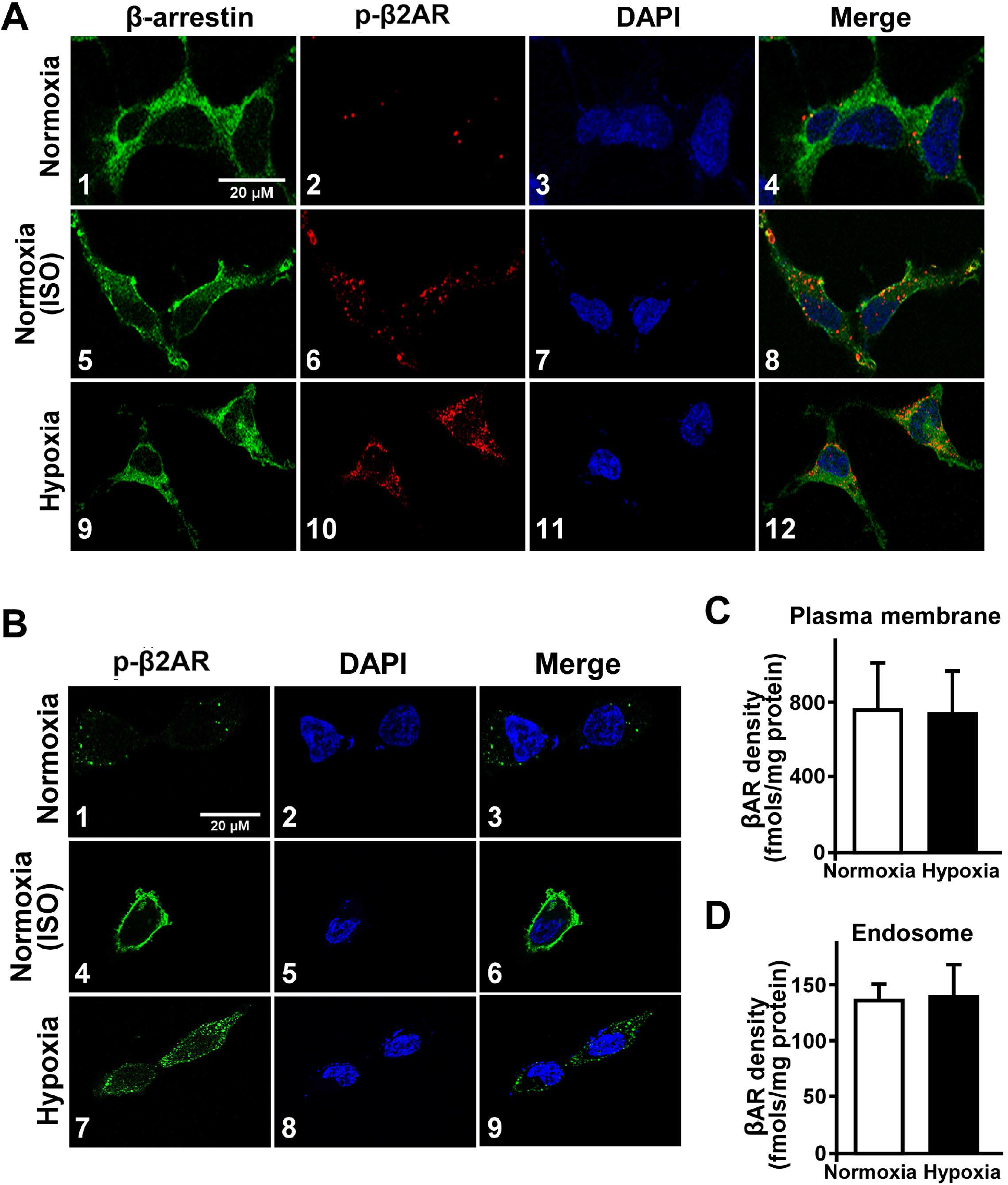
Hypoxia mediated non-canonical endosomal accumulation of phosphorylated β2ARs: **A**, Confocal microscopy was performed on β2AR-β-arrestin-2 GFP double stable HEK 293 cells following 6 hours of hypoxia in serum free media. β-arrestin-2 GFP was visualized by GFP (green) and phospho-β2AR by using anti-phospho-β2AR antibody (red). ISO (100 μM) stimulation for 10 minutes was used a positive control to show β-arrestin-2 GFP recruitment to plasma membrane. Nucleus was visualized by DAPI (blue) staining (n=3). Scale, 20 μm. **B**, β2AR-HEK 293 cells were pre-treated with internalization blockers (0.45 M sucrose and 2% β-cyclodextrin) and subjected to 6 hours of hypoxia or normoxia. Confocal microscopy was performed to visualize phospho-β2AR using anti-phospho-β2AR antibody (green) following hypoxia or 10 minutes of ISO treatment (used as positive control following 6 hours pre-treatment with internalization blockers) (n=3). Nucleus was visualized by DAPI (blue) staining. Scale, 20 μm. **C & D**, ^[125]^I-Cyanopindalol radio-ligand binding was performed on plasma membrane (C) or endosomal fraction (D) isolated from β2AR-HEK 293 cells following 6 hours of normoxia or hypoxia.

Since we observed marked accumulation of phosphorylated β2ARs in the cytosol following hypoxia, we tested whether hypoxia mediates internalization of receptors following β2AR phosphorylation by pretreating the cells with internalization blockers (sucrose and β-cyclodextrin [20]). Consistent with previous studies [20], ISO treatment resulted in significant phosphorylation of β2ARs (green) that decorates the plasma membrane as receptors do not internalize (n=3) [**Fig. 3B, panels 4 & 6**]. In contrast, despite pre-treatment with internalization blockers, accumulation of phosphorylated β2ARs were observed in the cytosol [**Fig. 3B, panels 7 & 9**] showing that internalization blockers do not alter hypoxia mediated phosphorylation and/or internalization. To directly test whether hypoxia mediates internalization of phosphorylated β2ARs, radio-ligand binding was performed on plasma membrane and endosomal fractions from β2AR-HEK 293 cells subjected to normoxia or hypoxia. Surprisingly, radio-ligand binding showed no appreciable differences in the β2AR distribution following hypoxia (n=6) [**Fig. 3C & D**]. These observations suggest that hypoxia mediates phosphorylation of endosomal β2ARs independent of internalization consistent with the findings internalization blockers [**Fig. 3B**].

### Endosomal accumulation of phosphorylated β2ARs with hypoxia is associated with inhibition of resensitization

Given the observation that hypoxia mediates β2AR phosphorylation independent of internalization, β2AR phosphorylation was assessed by immunoblotting of the plasma membrane and endosomal fractions following hypoxia. Significant β2AR phosphorylation was observed in the endosomal fractions of cells subjected to hypoxia when compared to normoxia while no appreciable differences were observed at the plasma membrane (n=5) [**Fig. 4A left and right panels**]. Endosomal β2ARs traditionally undergo dephosphorylation/resensitization by PP2A [26]. Since PP2A is acutely regulated by PI3Kγ activity [20], PI3Kγ was immunoprecipitated from plasma membrane and endosomal fractions and the immunoprecipitates were subjected to in vitro lipid kinase activity. Significant PI3Kγ activity was observed in the endosomal fractions following hypoxia compared to normoxia (n=4) [**Fig. 4B, right panel**] while no appreciable differences were observed in the plasma membrane fractions [**Fig. 4B, left panel**]. As endosomal PI3Kγ activity is higher, we assessed β2AR-associated phosphatase activity by immunoprecipitating β2ARs by using anti-FLAG antibody from plasma membrane and endosomal fractions. While no appreciable difference was observed in β2AR-associated phosphatase activity at the plasma membrane (n=6) [**Fig. 4C, left panel**], significant reduction in β2AR-associated phosphatase activity was observed in the endosomal fraction (n=6) [**Fig. 4C, right panel**]. Since we have previously shown that PI3Kγ inhibits PP2A activity by phosphorylating the endogenous inhibitor of PP2A, I2PP2A, immunoblotting was performed to assess I2PP2A phosphorylation using an in-house generated anti-phospho-I2PP2A antibody. Although total I2PP2A levels did not change [**Fig. 4D**], significant increase in I2PP2A phosphorylation was observed in hypoxia compared to normoxia (n=4) [**Fig. 4D, left and right panel**]. Despite reduced PP2A activity, there was no appreciable difference in the expression of PP2A following hypoxia [Fig. 4D]. These observations show that hypoxia mediates activation of PI3Kγ inhibiting resensitization leading to accumulation of phosphorylated β2ARs in the endosomal fractions accounting for receptor dysfunction.

**Figure 4.**
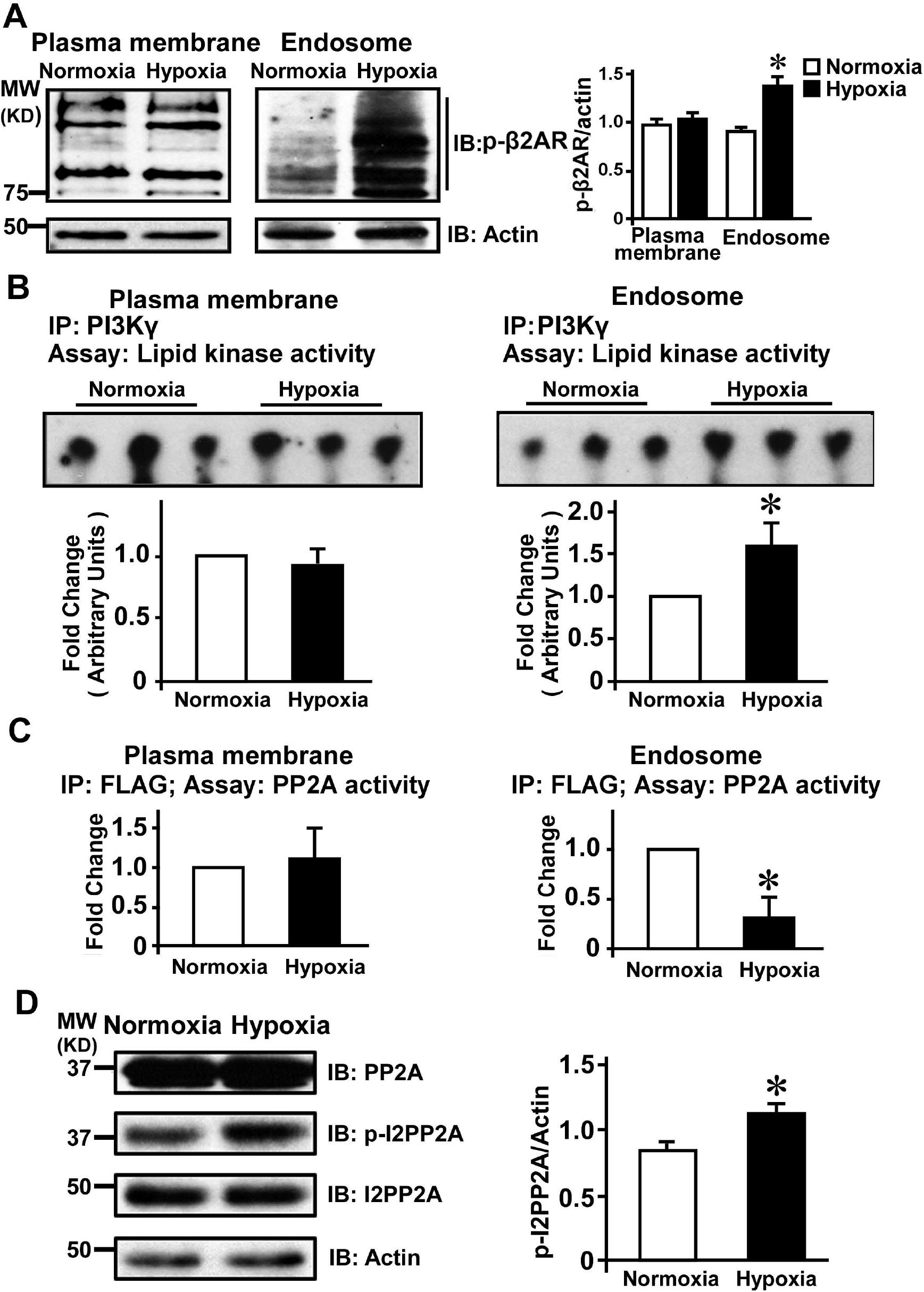
Hypoxia inhibits β2AR resensitization by impairing β2AR-associated PP2A activity: **A**, Plasma membrane (50 μg) or endosomal fractions (50 μg) from β2AR-HEK 293 cells were immunoblotted with anti-phospho-β2AR antibody following 6 hours of normoxia or hypoxia. Cumulative densitometry is shown as bar graphs (n=5). *p<0.05 vs normoxia. **B**, PI3Kγ was immunoprecipitated from plasma membrane or endosomal fractions (80 μg) from β2AR-HEK 293 cells following 6 hours of normoxia or hypoxia. The immunoprecipitated beads were washed and subjected to in vitro lipid kinase assay by providing phosphatidylinositol (PI) to generate phosphatidylinositol mono-phosphate (PIP) (upper panel) (n=4). Summary densitormetric data on generation of labeled PIP (lower panel). *p<0.05 vs normoxia. **C**, FLAG-β2AR was immunoprecipitated (IP) from plasma membrane (50 μg) or endosomal fractions (50 μg) and associated PP2A activity was measured in the FLAG immunoprecipitates (n=6). *p< 0.05 vs. normoxia. **D**, Western immunoblotting was performed on 80 μg total lysates from β2AR-HEK 293 cells following 6 hours of normoxia or hypoxia to detect PP2A, phospho-I2PP2A and I2PP2A. Actin was used as loading control (n=4). Cumulative densitometry is shown as bar graphs. *p<0.05 vs normoxia.

### Hypoxia causes adverse cardiac remodeling and is associated with β2AR dysfunction

Since it is known that hypoxia/ischemia is one of the leading causes of heart failure and stroke, studies were performed to assess whether acute hypoxia can cause deleterious cardiac remodeling. C57Bl6 mice were placed in hypoxia chamber for 20 hours [27] and cardiac function was assessed by echocardiography. Acute hypoxia resulted in deleterious cardiac remodeling as observed by increased cardiac lumen post-hypoxia (n=12) [**Fig. 5A, upper panel**] and as measured by functional parameters of % fractional shortening (%FS) and % ejection fraction (%EF) [**Fig. 5A, lower panel**]. Consistently, significant increase in heart weight to body ratio (HW/BW) was observed in mice subjected to hypoxia (n=12) [**Fig. 5B**] and H & E staining showed increased ventricular lumen following hypoxia (n=4) [**Fig. 5C**]. Since βARs are powerful regulators of cardiac function, we assessed whether acute hypoxia causes increase in cardiac β2AR phosphorylation. Immunoblotting of cardiac lysates showed significant increase in β2AR phosphorylation following hypoxia (n=6) [**Fig. 5D, upper panel and 5E, left panel**]. HIF-1α, the sentinel marker for hypoxia was also significantly stabilized in the hypoxia compared to normoxia [**Fig. 5D, middle panel and 5E, right panel**]. To test whether increased phosphorylation of β2AR is associated with receptor dysfunction, in vitro ISO-stimulated adenylyl cyclase activity was performed on the cardiac plasma membranes. There was significant reduction in adenylyl cyclase activity following hypoxia both at baseline and upon in vitro ISO stimulation (n=6) [**Fig. 5F**] which was preserved in normoxia. Together these findings show that acute hypoxia causes adverse cardiac remodeling and βAR dysfunction.

**Figure 5.**
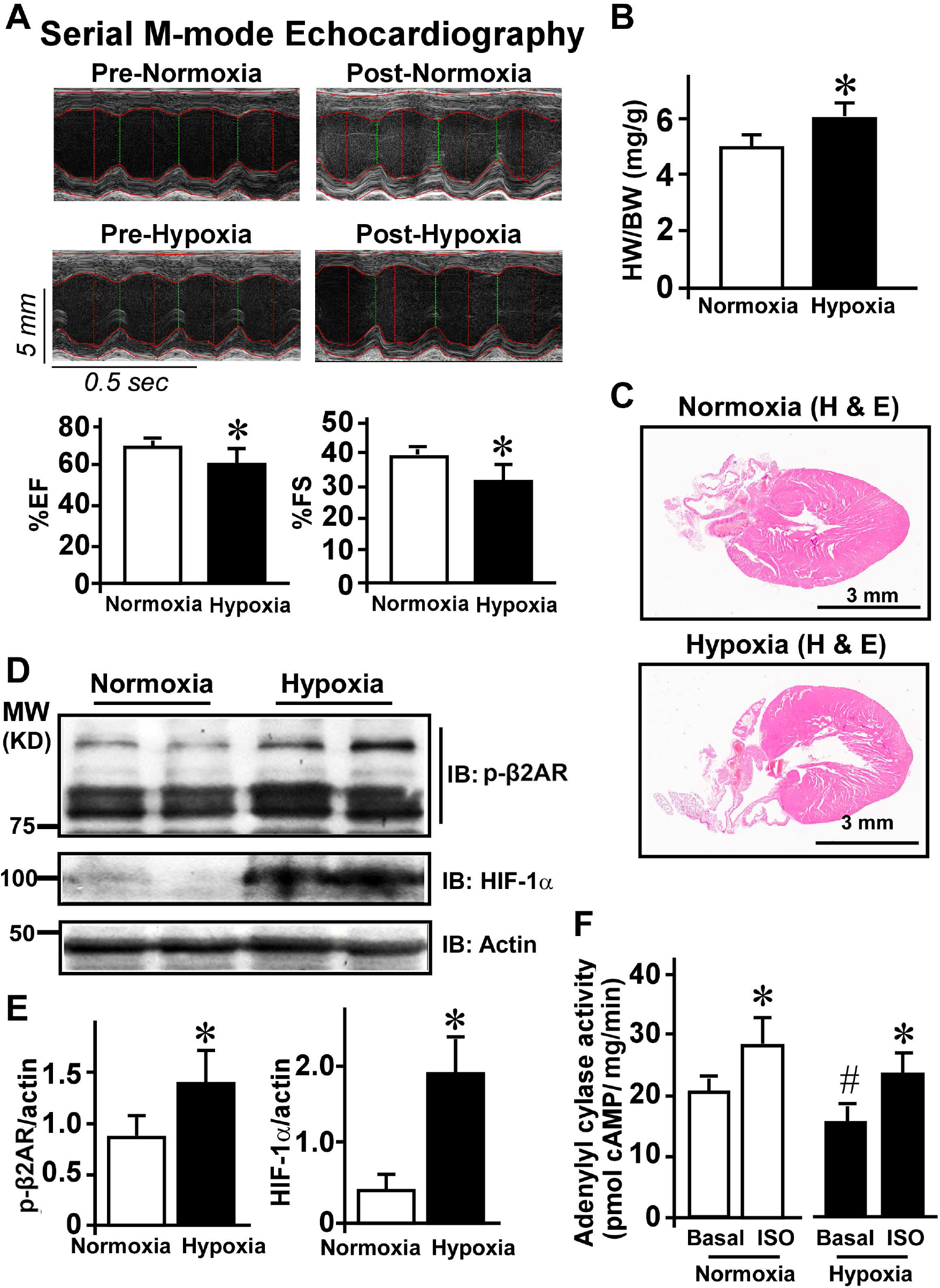
Acute hypobaric hypoxia in mice leads to adverse cardiac remodeling associated with βAR dysfunction: **A**, C57Bl6 mice were subjected to acute (20 hours) of hypobaric hypoxia. M-mode echocardiography was performed pre- and post-hypoxia or normoxia treatments. Acute hypoxia leads to larger ventricular chamber as assessed by echocardiography (n=12) (upper 4 panels). Lower panel shows cardiac functional parameters of % ejection fraction (%EF) (left panel, *p<0.05 vs. normoxia) and % fractional shortening (%FS) (right panel, *p<0.05 vs. normoxia) as measures of cardiac function. **B**, Heart weight (HW) and body weight (BW) were measured for the mice at the termination the experiment post-hypoxia or normoxia to assess HW/BW ratio as a measure of adverse cardiac remodeling (n=12). **C**, Heart sections from mice subjected to normoxia or hypoxia were stained with H & E to assess cardiac remodeling (n=4). H & E staining shows large ventricular lumen in mice subjected to hypoxia. Scale bar (3 mm). **D**, Total cardiac lysates (100 μg) were immunoblotted with anti-phospo-β2AR antibody (upper panel). The blots were stripped and re-probed with anti-HIF-1α a sentinel marker for hypoxic response (middle). Actin was used as a loading control. **E**, Cumulative densitometry data (n=6) for phospho-β2AR and HIF-1α is shown in the bar-graphs (left panel, *p<0.05 vs. phospho-β2AR normoxia; right panel, *p<0.01 vs. HIF-1α normoxia). **F**, In vitro isoproterenol (ISO)-stimulated adenylyl cyclase activity was measured in the cardiac plasma membranes isolated from the hearts of mice subjected to 20 hours of normoxia or hypoxia (n=6). *p<0.01 vs basal: #p< 0.05 vs. basal normoxia and ISO (normoxia or hypoxia).

### β-blocker reverses hypoxia-mediated β2AR phosphorylation

As β-blocker treatment in hypoxia reduces HIF-1α accumulation [9], experiments were conducted to test whether β-blocker pre-treatment alters the state of β2AR phosphorylation despite hypoxia. β2AR-HEK 293 cells were pre-treated with β-blocker propranolol followed by either hypoxia or normoxia and phosphorylation of β2AR was assessed by immunoblotting. Consistent with previous studies [28], significant phosphorylation of β2ARs was observed in normoxia in response to β-blocker (n=5) [**Fig. 6A & B**]. In contrast, β-blocker treatment in hypoxia surprisingly resulted in abrogation of hypoxia-mediated β2AR phosphorylation [**Fig. 6A & B**]. To further test whether β-blocker treatment results in loss of β2AR phosphorylation, confocal microscopy was performed following hypoxia. β-blocker treatment in normoxia resulted in marked increase of phosphorylated β2ARs as visualized by anti-phospho-β2AR antibody (green) (n=4) [**Fig. 6C & 6D (panels 5 and 6)**]. Hypoxia resulted in significant increase in phosphorylated β2ARs [**Fig. 6C & 6D (panels 3 and 4)**] consistent with our data [**Fig. 1**]. In contrast, β-blocker pre-treatment significantly reduced β2AR phosphorylation [**Fig. 6C & 6D (panels 7 and 8)**] showing a unique role of β-blocker in hypoxia. To further test whether unexpected reduction in phosphorylation by β-blocker in hypoxia is due to the ability of β-blocker to engage the resensitization pathway in hypoxia. Since hypoxia decreases endosomal β2AR-associated PP2A activity in hypoxia, FLAG-β2AR was immunoprecipitated from plasma membrane and endosomal fractions following hypoxia and β-blocker treatment to assess receptor associated activity. FLAG-β2AR associated PP2A activity was not appreciably different in the plasma membranes following β-blocker pre-treatment [**Fig. 6E, gray bar plasma membrane**]. However, there was significant increase in the FLAG-β2AR associated PP2A activity in the endosomes of cells pre-treated with β-blockers and subjected to hypoxia [**Fig. 6E, gray bar endosomes**]. This observation suggests that β-blockers may act differently in hypoxia than in normoxia wherein, they could mechanistically engage the resensitization pathway to reduce β2AR phosphorylation and underlie the benefits provided by β-blockers in patients with heart failure.

**Figure 6.**
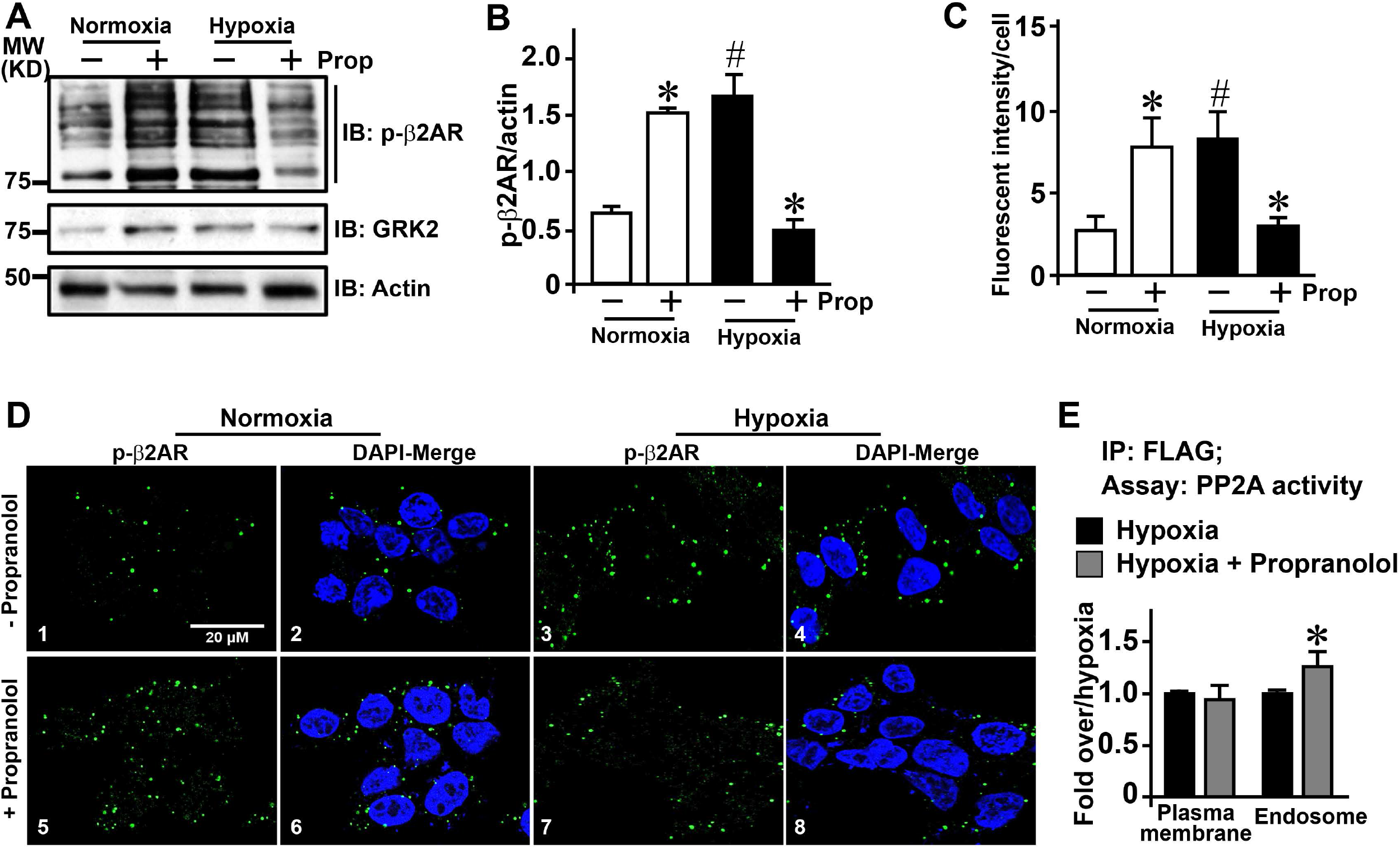
β-blocker reverses hypoxia-mediated β2AR phosphorylation: **A**, β2AR-HEK 293 cells were pretreated (45 minutes) with β-blocker propranolol (10 μM) and then subjected to normoxia or hypoxia for 6 hours. Total cell lysates (80 μg) were immunoblotted with anti-phospho-β2AR antibody to assess β2AR phosphorylation. The blot was stripped and re-probed with anti-GRK2 antibody and actin was used as loading control. **B**, Cumulative data (n=5) for phospho-β2AR is shown in the bar graph. *p<0.05 vs. respective vehicle (-propranolol); #p<0.05 vs. vehicle (propranolol) normoxia. **C**, Bar graph showing average fluorescent intensity/cell (n=4) (>100 cells/experiment). *p< 0.05 vs. respective vehicle (-propranolol); #p<0.05 vs. vehicle (- propranolol) normoxia. **D**, Confocal microscopy was performed to visualize phosphorylated β2ARs by using anti-phospho-β2AR antibody (green) and nucleus (blue) by DAPI after pretreatment with β-blocker propranolol (45 minutes) followed by 6 hours of hypoxia or normoxia. **E**, FLAG-β2AR was immunoprecipitated (IP) from plasma membrane (50 μg) or endosomal fractions (50 μg) following hypoxia alone or along with propranolol and associated PP2A activity was measured in the FLAG immunoprecipitates (n=8). *p< 0.05 vs. endosomal hypoxia.

## Discussion

Here we show that hypoxia leads to β2AR phosphorylation independent of its agonist indicating unique non-canonical regulation of the receptor. Hypoxia-induced β2AR phosphorylation is GRK2-dependent as GRK2 is selectively upregulated and GRK2 inhibition reverses β2AR phosphorylation. Hypoxia also leads to reduced cAMP and decreased adenylyl cyclase activity suggesting that GRK2 recruitment to the β2ARs may be independent of the Gβγ subunits [29, 30]. Although GRK2-mediates β2AR phosphorylation, there was no β-arrestin recruitment to plasma membrane or changes in dynamics of receptor internalization. Interestingly, hypoxia leads to selective accumulation of phosphorylated β2ARs in the endosomes with no changes at the plasma membrane. Importantly, hypoxia inhibits β2AR resensitization as β2AR-associated phosphatase activity was significantly impaired in the endosomes following hypoxia. Significant PI3Kγ activity was observed only in the endosomal fractions upon hypoxia consistent with inhibition of PP2A through the PI3Kγ-I2PP2A axis [20]. This is supported by the findings of significant I2PP2A phosphorylation showing that hypoxia mediates β2AR phosphorylation by activating GRK2 while simultaneously inhibiting PP2A which accounts for accumulation of phosphorylated receptors. These observations are strengthened by in vivo studies showing that acute hypoxia results in significant βAR dysfunction and is associated with adverse cardiac remodeling. β-blocker pre-treatment surprisingly reduced hypoxia-mediated β2AR phosphorylation and was associated with increased β2AR-associated phosphatase activity in contrast to its known role in mediating β2AR phosphorylation in normoxia.

Previous studies have shown that GRK2 phosphorylation of βARs is one of the key regulators of HIF-1α stabilization [9]. Consistently, our data shows that GRK2 is the key mediator of β2AR phosphorylation in hypoxia as inhibition of GRK2 results in loss of β2AR phosphorylation suggesting that GRK2 plays a critical role in hypoxia-mediated β2AR regulation. Traditionally, GRK2 is recruited to the βARs by the dissociated Gβγ subunits of the hetero-trimeric G protein following agonist stimulation of the receptor [11, 24]. However, hypoxia-mediated βAR phosphorylation is agonist independent suggesting that hypoxia may engage non-canonical pathways to mediate β2AR phosphorylation. In contrast to the classical GRK2-mediated phosphorylation that initiates receptor internalization through β-arrestin-dependent pathways [24, 31], there were no significant recruitment of β-arrestin to the β2ARs following hypoxia. Similarly, there were no differences in β2AR density/distribution between plasma membrane or endosomal fractions as assessed by radio-ligand binding studies. This suggests that hypoxia may not engage trafficking/internalization machinery to alter β2AR distribution despite our consistent observation of increased endosomal β2AR phosphorylation by confocal microscopy or western immunoblotting studies [**Figs. 1, 3 & 4**]. These set of unexpected observations brings forth a unique conceptual idea that hypoxia may directly initiate β2AR phosphorylation in the endosomes by GRK2 thus, by-passing the need of Gβγ subunits for recruitment. Such an idea would be consistent with phosphorylation and regulation of non-receptor substrates of GRK2 [30].

Classically following β2AR phosphorylation by GRKs, the receptor is endocytosed and resensitization occurs by dephosphorylation mediated by PP2A. Our previous study has shown that PI3Kγ regulates resensitization by inhibiting PP2A activity through phosphorylation of I2PP2A [20]. Also, agonist stimulation leads to kinase activation while, simultaneously inhibiting PP2A activity thus buttressing the kinase activation [11, 20]. Hypoxia leads to accumulation of phosphorylated β2ARs in the endosomes and is associated with significantly increased endosomal PI3Kγ activity and inhibition of PP2A activity. Furthermore, hypoxia leads to marked increase in phosphorylation of I2PP2A, the endogenous inhibitor of PP2A showing that mechanistically PP2A is inhibited by the PI3Kγ-I2PP2A axis. This shows that loss in PP2A activity and inability to dephosphorylate the receptor in part, contributes to the accumulation of phosphorylated β2ARs in the endosomes. PI3Kγ is also recruited to plasma membrane by the dissociated Gβγ subunits [32] but selective increase only in the endosomal PI3Kγ activity under hypoxia suggests non-canonical regulation of PI3Kγ. Hypoxia activation of PI3Kγ in the cytosol now mediates phosphorylation of I2PP2A inhibiting PP2A activity and thereby, dephosphorylation of the receptors. This consequently leads to impairment of resensitization accounting for accumulation of phosphorylated β2ARs in the endosomes.

Given the recognition that acute hypoxia could underlie stroke due to changes in cardiac function, mice were subjected to acute hypobaric hypoxia to assess cardiac remodeling. Interestingly, acute hypoxia resulted in adverse cardiac remodeling with left ventricular cardiac dysfunction associated with βAR dysfunction as assessed by adenylyl cyclase activity and β2AR phosphorylation. These observations suggest that βAR dysfunction in acute hypoxia may underlie the deleterious left ventricular cardiac remodeling compared to studies showing long term effects of hypoxia that were associated with marked alterations in the right ventricles [33, 34]. These studies support the idea that under conditions of acute hypoxia, the heart may have difficulty meeting the mechanical demands leading to stroke due to tissue hypoxia/ischemia. In this regard, recent studies have shown that in vivo β-blocker treatment markedly reduces renal HIF-1α stabilization and erythropoiesis [9]. While mechanisms underlying role of β-blockers in hypoxia are not well understood, our data surprisingly shows that β-blocker pre-treatment in hypoxia leads to loss in β2AR phosphorylation. This is in contrast to the observation that β-blockers mediate β2AR phosphorylation that initiates G protein-independent β-arrestin signaling in normoxia [35, 36]. Consistent with this paradigm, our data shows that β2ARs are significantly phosphorylated with β-blocker in normoxia but this phosphorylation is blocked in hypoxia due to the presence of the β-blocker. However, these unexpected observations may have significant clinical implications given that hypoxia *per se* initiates β2AR dysfunction through increased accumulation of phosphorylated receptors. Given that accumulation of phosphorylated βAR leads to reduced cardiac function/output [37–39] pre-disposing to the stroke, β-blocker treatment in these conditions may reverse the phosphorylation of βARs preserving cardiac function. This is supported by recent clinical trial wherein, use of β-blocker in patients with pulmonary hypertension (who are associated with HIF-1α glycolytic shift) showed increased cAMP [29]. This suggests that β-blocker in hypoxia may resensitize the β2ARs leading to increased cAMP that may potentially underlie the beneficial outcomes. Such an idea is supported by the observation of increased β2AR-associated phosphatase activity in the endosomes of cells pre-treated with β-blockers in hypoxia suggesting a yet to be understood role of β-blockers in hypoxia. Thus, our study shows that hypoxia mediates β2AR phosphorylation by selectively increasing GRK2-dependent kinase pathway and simultaneously inhibiting the PP2A phosphatase pathway through the PI3Kγ-I2PP2A axis [**Fig. 7**]. Also, unexpectedly our study identified that β-blocker reduces β2AR phosphorylation in hypoxia suggesting mechanisms beyond the current understanding of β-blocker function in normoxia and is being investigated.

**Figure 7.**
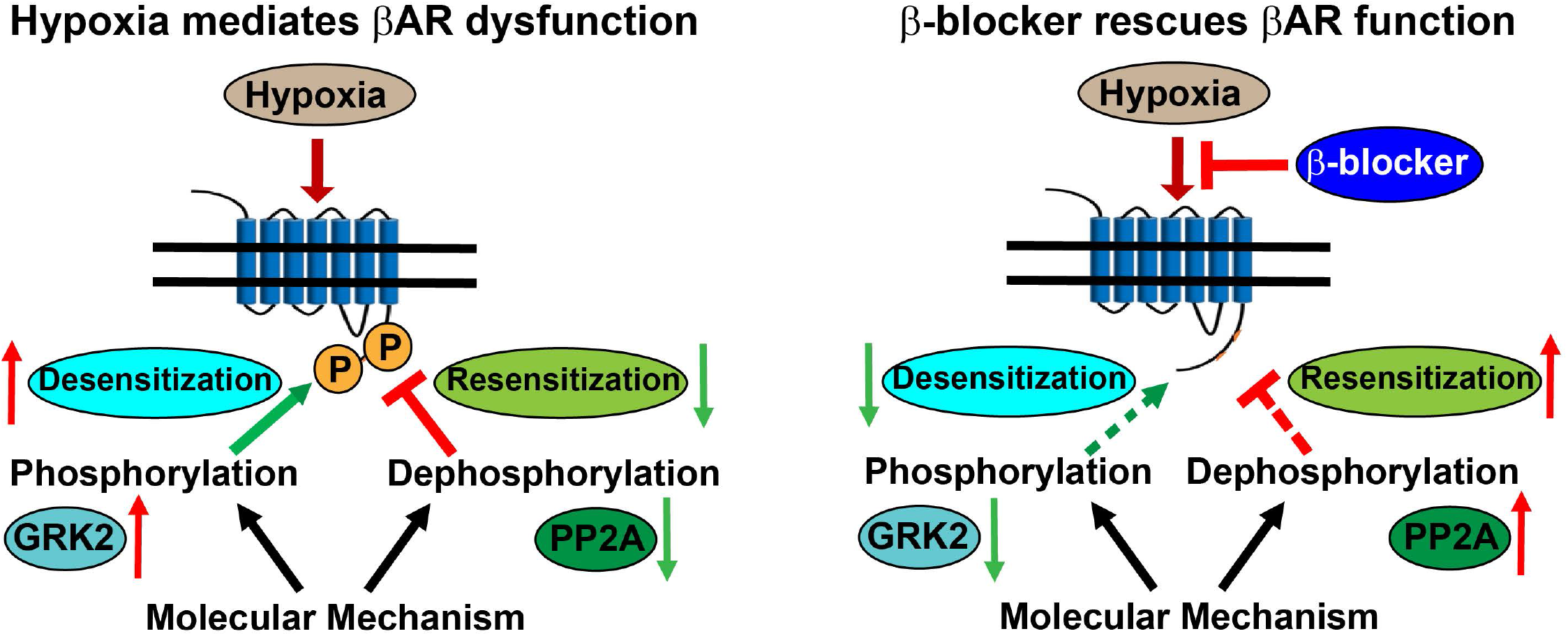
Schematic illustration: Proposed model showing that hypoxia non-canonically mediates β2AR dysfunction by selective upregulation of GRK2 that mediates receptor phosphorylation and endosomal accumulation of phosphorylated β2ARs. Simultaneously, hypoxia also impairs resensitization by inhibiting protein phosphatase 2A (PP2A) activity. Hypoxia inhibits PP2A by activating PI3Kγ that phosphorylates endogenous inhibitor of PP2A (I2PP2A) [20]. Phosphorylated-I2PP2A (phospho-I2PP2A) robustly binds to PP2A inhibiting PP2A activity [20]. Thus, the inability of PP2A to dephosphorylate β2ARs leads to impairment of resensitization and accumulation of phosphorylated in the endosomes. Surprisingly, β-blocker treatment reverses β2AR phosphorylation in hypoxia preserving receptor function by potentially reducing GRK2 levels and decreasing PI3Kγ activity normalizing PP2A that may now mediate β2AR dephosphorylation despite hypoxia. These studies bring-to-fore yet to be appreciated role of β-blockers in providing beneficial effects in hypoxia contrary to its currently understood role in normoxia.

## Methods

### Experimental Animal

C57/BL6 wild type (WT) mice of either sex 3-6 months of age were subjected to hypoxia (10% O2) [27] or normoxia for 20 hours. The studies were performed in accordance with institutional and national guidelines and regulations, as approved by Cleveland Clinic Institutional Animal Care and Use Committee.

### Cell Culture

HEK 293 cells were maintained in minimum essential media with 10% heat-inactivated fetal bovine serum and 1% penicillin/streptomycin [30]. Cells were seeded at a standard density of ~1-3 x 10^5^ cells /35 mm dish and the experiments were performed at 70-80% confluence. Cells were serum starved for 3 or 6 hours when treated with normoxia or hypoxia and for positive control cells were stimulated with 10μM isoproterenol (ISO) (Sigma-Aldrich) for 10 minutes in normoxia. For hypoxia studies, cells were incubated in a sealed chamber at 37°C with 2% O2, 5% CO2, balanced with 93% N2. GRK2 inhibitor Paroxetine (30nM) (Sigma-Aldrich) was added to β2AR-HEK 293 cells 45 minutes prior to normoxia or hypoxia for 6 hours. Similarly, the cells were pre-treated with propranolol (10 μM) (Sigma-Aldrich) for 45 minutes prior to incubation in hypoxia for 6 hours. β2AR//β-arrestin 2 GFP double stable cells were pre-treated with endocytosis inhibitors 0.45M sucrose and 2% β-cyclodextrin for 1 hour and subjected to normoxia or hypoxia treatment. For control ISO treatment, the cells were pre-treated with endocytosis blocker for 1 hours and stimulated with 10 μM ISO for 10 minutes. HEK 293 cells stably expression FLAG-β2AR (β2AR-HEK 293) cells was a generous gift from R. Lefkowtiz, Duke University, Durham, NC. HEK293 cells stably overexpressing β2AR and β-arrestin 2 GFP was a generous gift from Dr. Marc G. Caron, Duke University, Durham, NC.

### Isolation of Plasma Membranes and Early Endosomes

Plasma membranes and early endosomes were isolated as previously described [20]. Plasma membranes were prepared by homogenizing of samples in non-detergent lysis buffer (5mmol/L Tris-HCl pH 7.5, 5 mM EDTA, 1 mM PMSF, and 2 μg/mL Leupeptin and Aprotinin). Cell debris/nuclei were removed by centrifugation at 1000 x *g* for 5 minutes and the supernatant was centrifuged at 30,000 x *g* for 30 minutes. Pellet representing membrane fraction was suspended in 75 mM Tris-HCl pH 7.5, 2 mM EDTA, and 12.5 mM MgCl_2_ while supernatant was centrifuged for 1 hour at 100,000 rpm to obtain early endosomes. Endosomes as pellets were resuspended in the same buffer as used on plasma membranes.

### Confocal microscopy

β2AR-HEK 293 cells were plated onto glass coverslips pre-treated with 0.01% poly L-Lysine (Sigma-Aldrich). Cells were serum starved while cultured in normoxia or hypoxia incubator for 6 hours or stimulated with ISO in normoxia as positive control along with endocytosis inhibitors. The cells were fixed with 4% paraformaldehyde, permeabilized with 0.1% Triton X-100, and incubated in 5% goat serum in PBS. Anti-phospho-β2AR 355/356 [40] antibody (1:500, Santa cruz) diluted in 1% goat serum was used as primary antibody, and anti-rabbit Alexa Flour 488 (1:500) was used a secondary antibody (Molecular Probes). Similar treatments were performed for HEK293 cells stably overexpressing β2AR and β-arrestin 2 GFP except that phospho-β2AR were visualized by using anti-rabbit Alexa Flour 568 (1:500) as secondary antibody (Molecular Probes). Samples were visualized using sequential line excitation at 488 and 568 nm for green and red, respectively. 70 to 100 positive cells were analyzed in each experiment and quantitation was performed using IMAGE PRO PLUS7 (Media Cybernetics, Inc).

### Western immunoblotting

Standard procedure for western immunoblotting were performed as described previously [30]. The proteins were resolved on SDS-PAGE and transferred to PVDF (BIO-RAD) and assessed for protein using primary anti-bodies as described below. Antibodies for HIF-1α (1:500), phosphorylated-β2AR (1:1000), PI3Kγ (1:200), I2PP2A (1:5000), GRK2, 3, 5, 6 diluted at 1:1000 were from Santa Cruz Biotechnology, Flag antibody (1:1000) was from Roche, PP2Ac antibody (1:2000) was from Upstate Biotechnology (Millipore), β-actin antibody (1:20000) was from Sigma. Antibody for phosphorylated I2PP2A (anti-phospho-I2PP2A) (1:1000) was generated in house and described in our recent publication [41].

### Phosphatase assay

PP2A phosphatase activity was measured using phosphatase assay kit (Upstate Biotechnology, Millipore) following manufacturer’s protocol. Immunoprecipitated samples were resuspended in the phosphate free assay buffer and incubated in presence or absence of PP2A specific Serine-Threonine phospho-peptide substrate for 10 minutes. The reaction mix was incubated with acidic malachite green solution and absorbance was measured at 630 nm in a plate reader.

### Lipid kinase assays

Assays were performed as previously described [42]. Briefly, cells were solubilized in Triton X100 lysis buffer (0.8% Triton X-100, 20 mM Tris-HCl pH 7.4, 300 mM NaCl, 1 mM EDTA, 20% glycerol, 1 mM PMSF, 2 μg/ml each of Leupeptin and Aprotinin). PI3Kγ was immunoprecipitated using anti-PI3Kγ antibody. The beads were washed with lysis buffer and suspended in reaction buffer TNE (10 mM Tris-HCl, pH 7.4, 150 mM NaCl, 5 mM EDTA, and 100 μM sodium-orthovandate,). To the resuspended beads, 10 μl of 100 mM MgCl_2_, 10 μl of 2 mg/ml PtdIns (20 μg) sonicated in TE buffer (10 mM Tris–HCl pH 7.4 and 1 mM EDTA), 10 μl of 440 μM ATP containing 10 μCi of ^32^P-γ-ATP were added. The assay was performed at 23°C for 10 min with continuous agitation and stopped by 6N HCl. Lipids were extracted by chloroform:methanol (1:1) and spotted on to 200 μm silica-coated TLC plates (Selecto-flexible; Fischer Scientific, Pittsburgh, PA), and phosphorylation was assessed by autoradiography.

### βAR Density, Adenylyl Cyclase Activity and cAMP assays

βAR density was measured as described previously [32]. Briefly, 20 μg plasma membranes or endosomes were incubated with 250 pmol of ^[125]^I-Cyanopindolol alone or along with 40 μmol/L ICI (to determine nonspecific binding) at 37°C for 1 hour. The non-specific counts in presence of ICI were subtracted from the total ^[125]^I-Cyanopindolol counts to calculate for the receptor density. Adenylyl cyclase assays were determined by incubating 20 μg of membranes or endosomes at 37°C for 15 min with vehicle, isoproterenol or NaF (G-protein activator) in 50 μL of assay mixture containing 20 mM Tris-HCl, 0.8 mM MgCl_2_, 2 mM EDTA, 0.12mM ATP, 0.05 mM GTP, 0.1 mM cAMP, 2.7 mM phosphoenolpyruvate, 0.05 IU/ml myokinase, 0.01 IU/ml pyruvate kinase and ^32^P-α-ATP and generated cAMP was quantified by scintillation counting [30]. The cAMP content in the lysates was determined according to the manufacturer’s instruction by catch point cAMP immunoassay kit (Molecular Devices) [20].

### Immunohistochemistry

Freshly harvested cardiac samples were placed in fresh 4% paraformaldehyde at room temperature for 24 hours, followed by ethanol dehydration, xylene exchange, wax soaking and embedding tissues into paraffin blocks. Paraffin slides (5 μm thickness) were subsequently stained with H&E. Photographs were taken using a Slide Scanner-Aperio AT2 (Leica Biosystems).

### Echocardiography

Echocardiography was performed on anesthetized 8 −12 weeks old mice using a VEVO 2100 (VisualSonics) pre- and post-hypoxia treatment as previously described [43]. The mice in normoxia also were imaged at the same time. M-mode views were recorded including left ventricular systolic and diastolic dimensions, septum, and posterior wall which were used to calculate the functional parameters.

### Statistical analysis

Results are expressed as means ± SD. Data were analyzed by *t test* for two-group comparison (for example, βAR density in plasma membrane or endosome in Fig. 3B). For comparison of more than two groups, we used one-way analysis of variance (ANOVA) if there was one independent variable (for example, p-β2AR densitometric analysis in Fig. 1A) and two-way ANOVA if there were two independent variables (for example, p-β2AR densitometric analysis in Fig. 2B and adenylyl cyclase assay in Fig. 5E). A probability value of <0.05 was considered significant.

## Acknowledgments

We would like to thank Dr Sadashiva Karnik and Dr. Sarah Schumacher-Bass for constant feedback and insightful thoughts during the development and progression of the project. Funding: These studies were supported by NIH R01 HL089473 and HL128382 (SVNP).

